# Development of a model estimating root length density from root impacts on a soil profile in pearl millet (*Pennisetum glaucum* (L.) R. Br). Application to measure root system response to water stress in field conditions

**DOI:** 10.1101/574285

**Authors:** A. Faye, B. Sine, J.L. Chopart, A. Grondin, M. Lucas, A. Diedhiou, P. Gantet, L. Cournac, D. Min, A. Audebert, A. Kane, L. Laplaze

## Abstract

Pearl millet, unlike other cereals, is able to withstand dry and hot conditions and plays an important role for food security in arid and semi-arid areas of Africa and India. However, low soil fertility and drought constrain pearl millet yield. One of the main targets to address these constraints through agricultural practices or breeding is root system architecture. In this study, in order to easily phenotype the root system in field conditions, we developed a model to predict root length density (RLD) of pearl millet plants from root intersection densities (RID) counted on a trench profile in field conditions. We identified root orientation as an important parameter to improve the relationship between RID and RLD. Root orientation was notably found to differ between thick roots (more anisotropic with depth) and fine roots (isotropic at all depths). We used our model to study pearl millet root system response to drought and showed that pearl millet reorients its root growth toward deeper soil layers that retain more water in these conditions. Overall, this model opens ways for the characterization of the impact of environmental factors and management practices on pearl millet root system development.

## Introduction

Pearl millet (*Pennisetum glaucum* (L.) R. Br., syn. *Cenchrus americanus* (L.) Morrone) is a cereal crop domesticated in the Western part of Sahel about 5,000 years ago [1]. It is well adapted to dry tropical climate and low-fertility soils and therefore plays an important role for food security in arid and semi-arid regions of sub-Saharan Africa and India. In these areas, pearl millet is one of the most important sources of nutritious food [2, 3] and is the staple crop for nearly 100 million people [4, 1]. Its grain is rich in protein, essential micronutrients and calories. It is also gluten-free and has hypoallergenic properties [4]. In a context of climate change leading to unpredictable weather patterns and rising temperatures in West Africa [5, 6], pearl millet could play an even more important role for food security because it can withstand hot and dry conditions that would lead to the failure of other locally grown cereal crops such as maize or sorghum. However, pearl millet lags far behind other cereals in terms of breeding and its yield is low. The recent sequencing of a reference genome and about 1, 000 accessions [4] open the way for a new era of genomic-based breeding in pearl millet. However, this will depend on the availability of phenotyping methods to characterize and exploit the available genetic diversity and identify interesting target traits.

Drought and low soil fertility are among the most important factors limiting pearl millet yield. The root system is responsible for water and nutrient uptake, and root system architecture is therefore a potential target in pearl millet breeding program to address these constraints. It is also an important trait to consider when analyzing the impact of agricultural practices. However, despite tremendous progress in the genetic characterization of root development, root system architecture phenotyping remains challenging particularly in agronomically-relevant field conditions. The root length density (total length of roots per unit of soil volume; RLD) is a key factor to estimate the soil volume explored by a root system and consequently the amount of water and nutrients available to the plant [7–12]. Therefore, RLD could be used to screen drought-resistant varieties.

The aim of this study was to develop a technique to map RLD in pearl millet plants from simple measurements in field conditions. For this we analyzed the relationship between RLD and root intersection densities (number of roots intersecting a vertical plane per unit of surface; RID) counted on trench profiles in field conditions. From this, we computed and experimentally validated a simple mathematical model linking RLD to RID. We then used this model to study the effect of drought stress on pearl millet root system architecture in two pearl millet varieties.

## Materials and methods

### Plant material

Four millet varieties were used for model calibration (Exp. 1): Souna3, Gawane, Thialack2 and SL87 (Table 1). Six varieties were tested for model validation (Exp. 2): Souna3 (common between Exp. 1 and Exp. 2), IBV8004, GB8735, ISMI9507, SL423, and SL28 (Table 1). The impact of water stress on pearl millet root system development was tested in a third experiment (Exp. 3) on SL28 (dual-purpose variety) and LCICMB1 (inbred line; [13]).

**Table 1:**
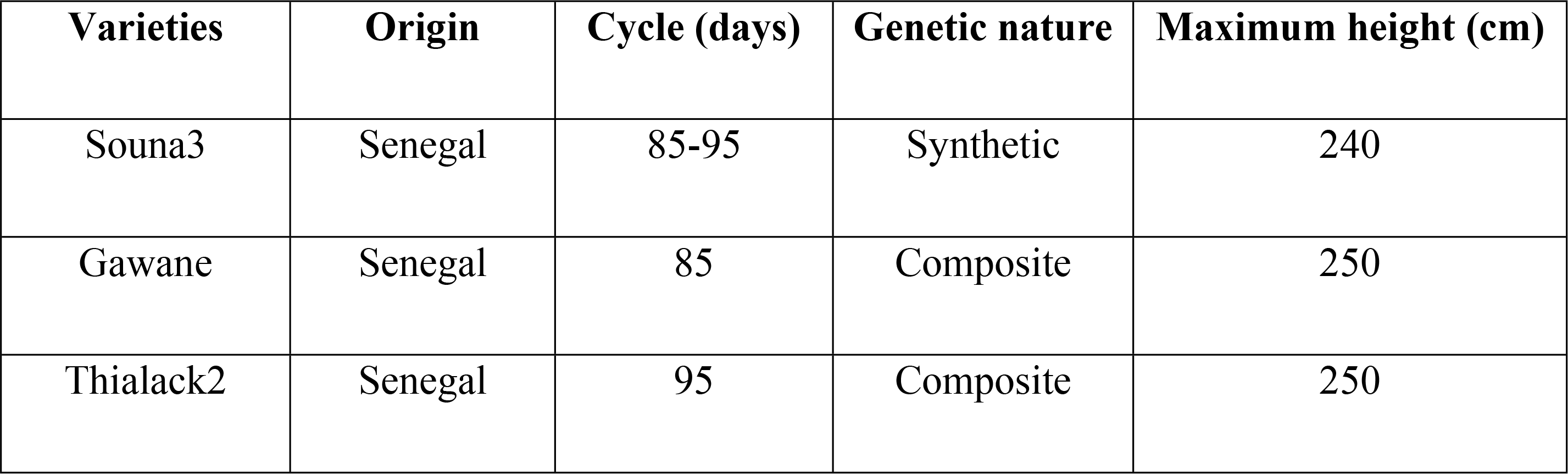
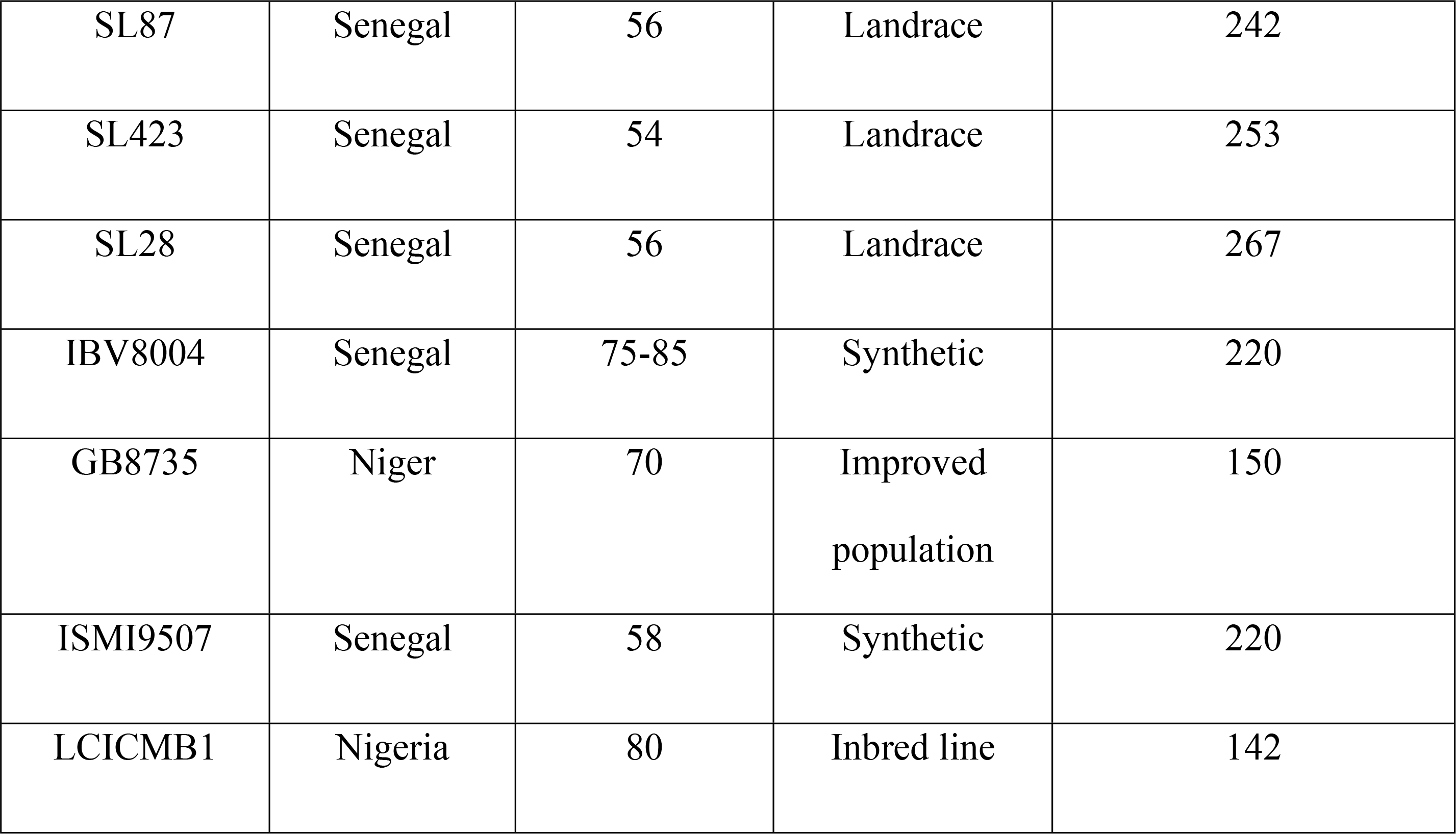
Pearl millet varieties used in this study.

| Varieties | Origin | Cycle (days) | Genetic nature | Maximum height (cm) |
| --- | --- | --- | --- | --- |
| Souna3 | Senegal | 85-95 | Synthetic | 240 |
| Gawane | Senegal | 85 | Composite | 250 |
| Thialack2 | Senegal | 95 | Composite | 250 |
| SL87 | Senegal | 56 | Landrace | 242 |
| SL423 | Senegal | 54 | Landrace | 253 |
| SL28 | Senegal | 56 | Landrace | 267 |
| IBV8004 | Senegal | 75-85 | Synthetic | 220 |
| GB8735 | Niger | 70 | Improved<br>population | 150 |
| ISMI9507 | Senegal | 58 | Synthetic | 220 |
| LCICMB1 | Nigeria | 80 | Inbred line | 142 |

### Site characteristic and experimentations

Field trials were performed at the Centre National de Recherche Agronomique station (CNRA) of the Institut Sénégalais des Recherches Agricoles (ISRA) in Bambey, Senegal (14.42°N, 16.28°W, altitude 17 m) in collaboration with and with the permission of the ISRA. Trials did not involve endangered or protected species. Exp. 1 was performed for model calibration during the rainy season 2016, Exp. 2 was performed in the dry season 2017 for model validation and Exp. 3 was performed in the dry season 2018 for response of pearl millet to a water stress. Exp. 2 and 3 were performed in the dry season in order to fully control the irrigation regime. Soil in the field trials was sandy and had the typical characteristics of the West Africa Sahelian soils in which pearl millet is grown. Tillage and chemical fertilization were applied as recommended for pearl millet [14]. Weeding was performed before planting.

Exp. 1 and Exp. 2 were laid out in a randomized complete block design with four plots per variety, each with five rows of 4 m long with a spacing of 0.8 m between plants and rows. In Exp. 1, water was provided by rainfall and additional irrigation was provided when needed (S1 Fig). Water stress was quantified using the PROBE water balance model [15]. The water balance simulations showed a decrease in the daily actual evapotranspiration to maximum evapotranspiration (AET/MET) ratio at the end of the cycle (S1 Fig). In Exp. 2, field was irrigated twice a week until 70 days after planting (DAP) and rainfall occurred at the end of the cycle (S1 Fig). The AET/MET ratio decreased only during the last days of cropping cycle (S1 Fig).

Exp. 3 was laid out in a randomized complete blocks design with split-plot into four blocks, the whole plots were for the water regime and the split-plots were for the varieties treatments. Plots consisted in four rows of 4 m long with 0.80 m between plants and rows. Thinning was done on eight days after emergence, at the rate of 2 plants per planting hole. In the well-watered plots (WW), irrigation was performed twice per week with 30 mm water per irrigation. In the drought stress plots (DS), a water stress was applied by withholding irrigation from 40 DAP for 32 days (S2 Fig) leading to a strong decrease in the AET/MET ratio (S2 Fig). At 72 DAP irrigation was resumed until the end of the growth cycle in addition with the first rain in June (S2 Fig). Field dry-down was monitored by measuring volumetric soil moisture to evaluate the fraction of transpirable soil water (FTSW) using Diviner probes (Sentek Pty Ltd) as previously described [14].

### Roots sampling for model development

We adapted a method previously described to estimate the RLD from intersections between roots and the face of a soil trench profile (root intersection density or RID; [10–12, 16 and 17]). Trench profiles were dug perpendicularly to the sowing rows and at two distances (30 then 10 cm) from the plant stalk base (Fig 1A). Three-sided incomplete steel cubes with sharpened edges facilitating penetration into the soil were used to sample soil cubes (Fig 1BC). The sampling device was pressed into open soil profile (trench profile) until its rear plane was aligned with the soil profile (Fig 1D) and then cut out of the soil to obtain a cube of soil (Fig 1E). A second sample was taken at the same depth and distance from the plant but with the open sides oriented in the opposite direction, in order to have open soil planes for six sides of the cube. Sampling was made at six depth levels ranging from 0.1 to 1.1 m and at two different dates (60 and 80 days after sowing, DAS). For each soil cube, the number of impacts (number of roots intersecting a plane, NI) on each side (transversal, longitudinal and horizontal; Fig 1B) was counted in the field immediately after sampling (Fig 1F; [10–12]). Thereafter, roots were washed out of the sampled soil cubes using a sieving conventional technique. Root lengths were measured for thick (d >1 mm) and fine roots (d < 1 mm) after scanning and analysis with WinRhizo (v 4, Regent Instruments, Inc, Quebec, Canada). Measurements were repeated four times per variety (384 cubes in total) and repeated measurements were averaged (i.e., same variety, same seeding rate, same date and same position).

**Fig 1.**
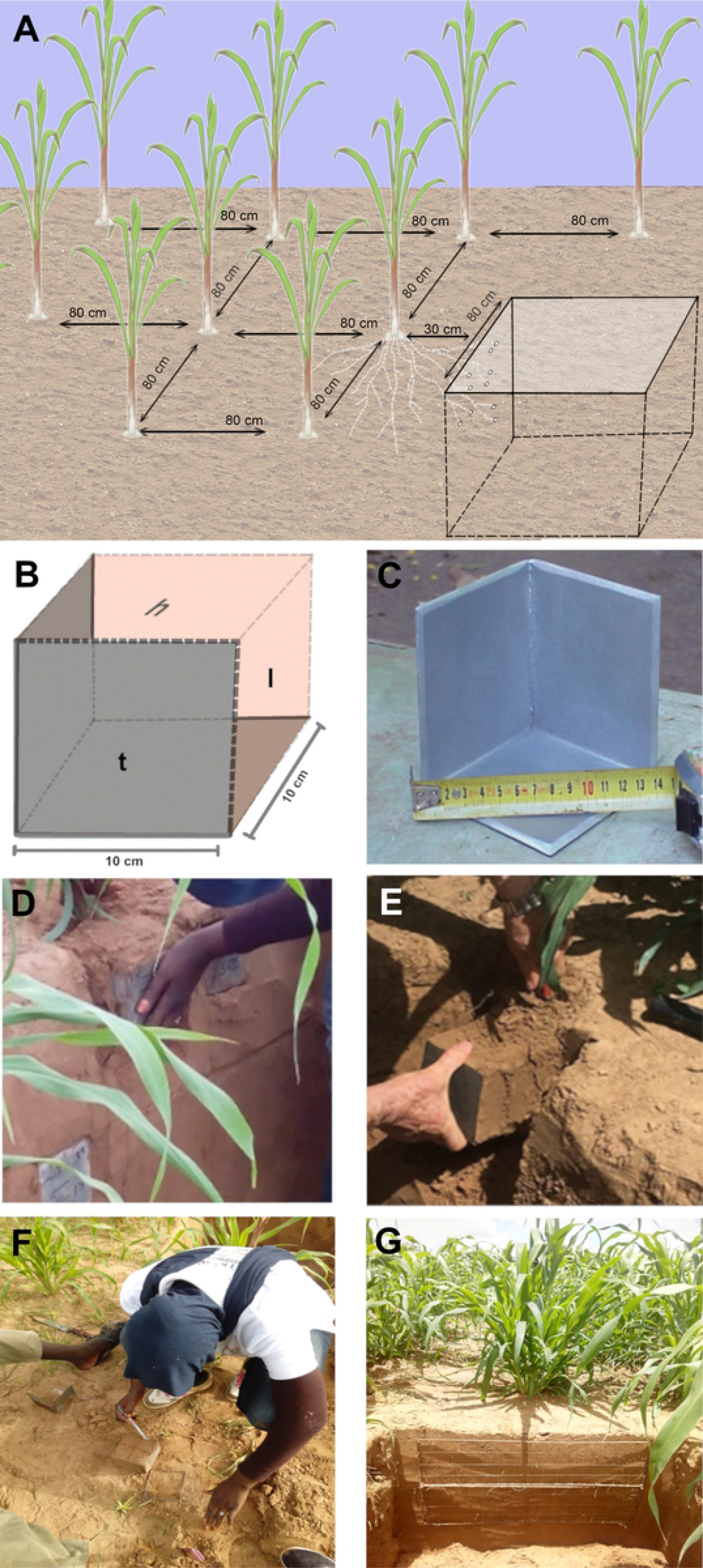
Root intersections density (RID) counting method used for root length density (RLD) modeling from RID. (A) Experimental design and trench profile for root sampling (at 30 cm from the plant in this example), (B) and (C) sampling device with sides oriented according to the soil surface and plant row (H: horizontal, L: longitudinal, T: transversal), (D) and (E) root sampling process, (F) Root impacts counting on all three sides of soil cubes extracted from the trench profile, and (G) grid (5x 5 cm mesh) on a soil profile for soil-roots intersections counting (RI).

The same protocol was used in the validation test, except that measurements were performed at four sampling dates (21, 40, 60 and 80 DAP) at eight soil depths ranging from 0.1 to 1.6 m. Measurements were repeated three times per variety (725 cubes in total). Soil samples containing less than three roots on one side of the cube were not considered.

### Model construction and test

A model was developed to establish a relationship between root impacts (root intersection densities, RID) counted on the two vertical planes of the cubes (longitudinal, *l*, and transversal plane, *t*) and the measured root length density (RLD_m_) contained in the soil samples collected. RLD of fine and thick roots were calculated (RLD_c_) on the basis of RID measured in a vertical soil plane using a direct empirical relationship first, and then considering the root distribution (anisotropy, root preferential orientation (P) as proposed by Lang and Melhuish [18]). A vertical index (*P*v) was calculated for the two vertical planes (*l* and *t*) using root counted on three faces of a soil cube (*l, t* and horizontal, *h*) as follow:

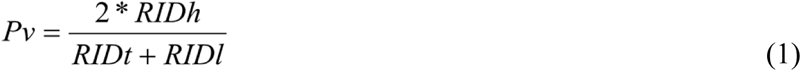

If P_v_ > 1 or < 1, the roots have a parallel or perpendicular preferential orientation with respect to the reference plane v. Depending on whether the Pv is =, > or < to 1, three RLD equations can be considered to calculate RLDc from RIDv of all Pv values [10, 19, and 20]. They can be combined in a general relationship using a synthetic root orientation coefficient (CO) dependent on Pv index values as described in Equation (3):

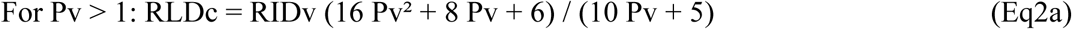

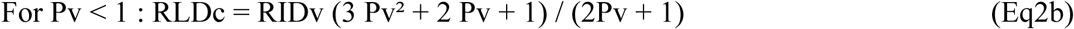

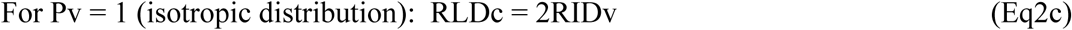

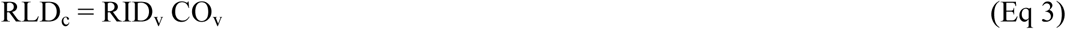

### Mapping of root intersections density (RID) on a trench-profile

In order to test the impact of water stress on pearl millet root development, a trench-profile method was used [20, and 21]. Soil-root intersections on a trench profile were counted using a 5 cm mesh grid applied to the soil profile (Fig 1G). Root intersections were counted at two distances (30 then 10 cm) from the base of the stalks and until no more roots could be found on the vertical dimension of the trench-profile. Root counting was performed at two dates of the cropping cycle: at the beginning of a stress (44 DAP) and at the end of stress (72 DAP). At each date, four soil profiles were measured per variety.

### Statistical analyses

Excel 2013 (Microsoft Corporation) was used for data cleansing and synthesis, to calculate anisotropy and preferential orientation indexes and to develop and test the obtained models. SPSS and R softwares (IBM Corp. Released 2016. IBM SPSS Statistics for Windows, Version 24.0. Armonk, NY: IBM Corp and R Development Core Team (2008). URL http://www.R-project.org.) were used to study the relationships linking the direction indexes and the experimental factors through an analysis of variance and a Student’s independence test at the 5% threshold. For model development study, the quality of the relationships between the RLD values measured in soil cubes (RLD_m_) and those modelled (RLD_c_) were evaluated taking into account slope, intercept and regression (R^2^). Nash’s Efficiency Ratio (NE; [22]), root mean square error (RMSE; [23]) and mean bias were used to compare measured (RLD_m_) and calculated (RLD_c_) deviations.

## Results

### Development of a model to extrapolate RLD from RID in field conditions

Four pearl millet varieties were selected to measure root intersections density (RID) and root length densities (RLD_m_) and try to create a model estimating root length densities in field conditions. We first analyzed the diversity of these four pearl millet varieties for root and shoot characters. There were significant differences in root length densities (RLD_m_) and root biomass densities (Fig 2A,B). Some varieties had deeper root systems than others. By contrast no significant differences were observed between varieties for shoot traits such as biomass (Fig 2C). Hence, these four varieties had contrasted root systems and were deemed suitable for model development.

**Fig 2.**
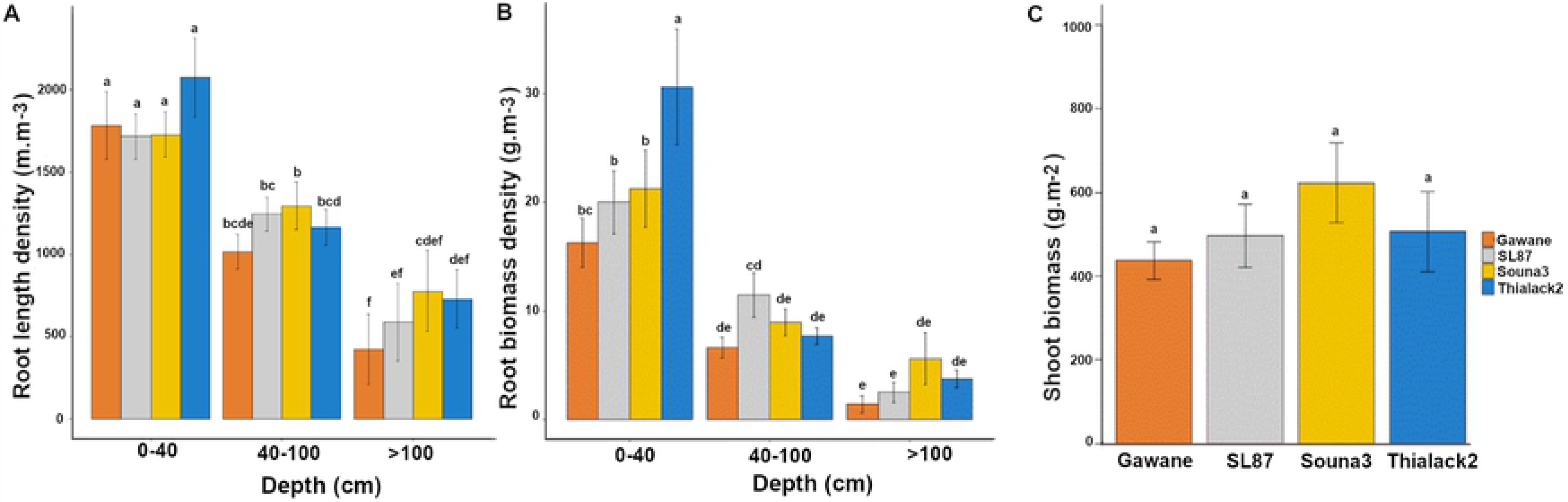
Characteristics of the varieties used for model calibration. **(A)** Root length density, (B) root biomass density and (C) shoot biomass. Data are mean +/− standard deviation. Significant differences (Tukey’s HSD) are indicated by different letters. For root traits, the mean of the two observation dates (60 DAP and 80 DAP) was considered.

We then analyzed the relationships between measured RLD and root intersection densities (i.e. the number of root impacts on a soil surface per surface unit; RID) on the 3 sides of a soil cube. Cubic soil samples were taken at 30 then 10 cm from the plant at different depth at 60 and 80 DAP. The number of root intersections (NI) was counted on 3 sides (vertical, transversal and horizontal) of the cube and then roots were washed out of the sampled soil cubes using a sieving conventional technique and root length was measured to compute a measure RLD (RLD_m_). Two classes of roots (thick >1 mm and fine < 1 mm) were considered. The simple linear regression between the number of impacts (NI) counted for all roots in a vertical plane and RLD_m_ for all roots showed unsatisfactory fit (RLD= 1.83 NI; R^2^ = 0.575, n= 70; Fig 3) indicating that more parameters needed to be included.

**Fig 3.**
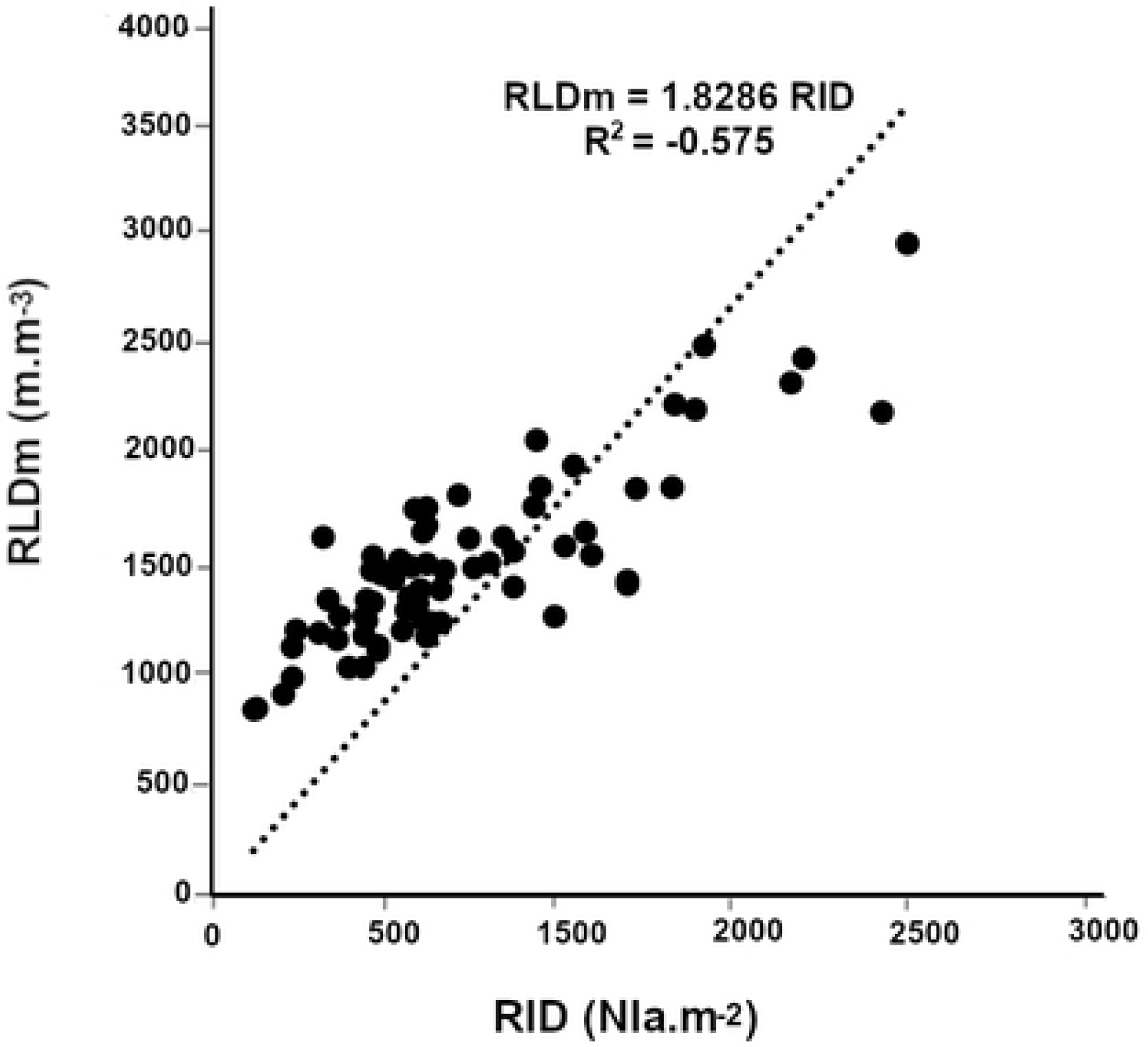
Fine and thick root growth orientation. Relationship between the number of measured root impacts on a vertical face and measured root length density.

Previous studies on several crops revealed different growth orientations for roots depending on soil depth [10, 11]. We therefore analyzed the impact of soil depth (seven soil depths between 10 and 130 cm), plant varieties and distance to the plant stalk base on root growth orientation. The root preferential orientation (Pv) was estimated from three-sided counts of a cube and used to calculate a root orientation coefficient (COv). We observed that the main root growth direction estimated by the Pv coefficient were not significantly different between varieties, measuring dates (60 JAS and 80 JAS) or sampling distances (10 or 30 cm) to the plant (S1 Table). The Pv index only depended on depth. As a consequence, the results from all varieties, measurement dates and sampling distances were pooled and we only analyzed the relationship between depth and root growth orientation. Considering all roots, we found a linear relationship between the root orientation index on a vertical plane (P) and depth (Z in meters; Fig 4A; Pv = 0.3408 Z + 0.905; R^2^ = 0.843, n= 70). Similarly, the root orientation coefficient (COv) was closely dependent on root depth (Fig 4B; B; COv = 0.471 Z +1.869; R^2^= 0.839, n= 70). COv values ranged from 1.92 at 0.10 m depth to 2.44 at 1.4 m depth. It was close to 2 close to the surface (0.10 m), indicating that roots had no preferential growth direction in the topsoil layers and that they gradually grew more in a vertical direction with depth.

**Fig 4.**
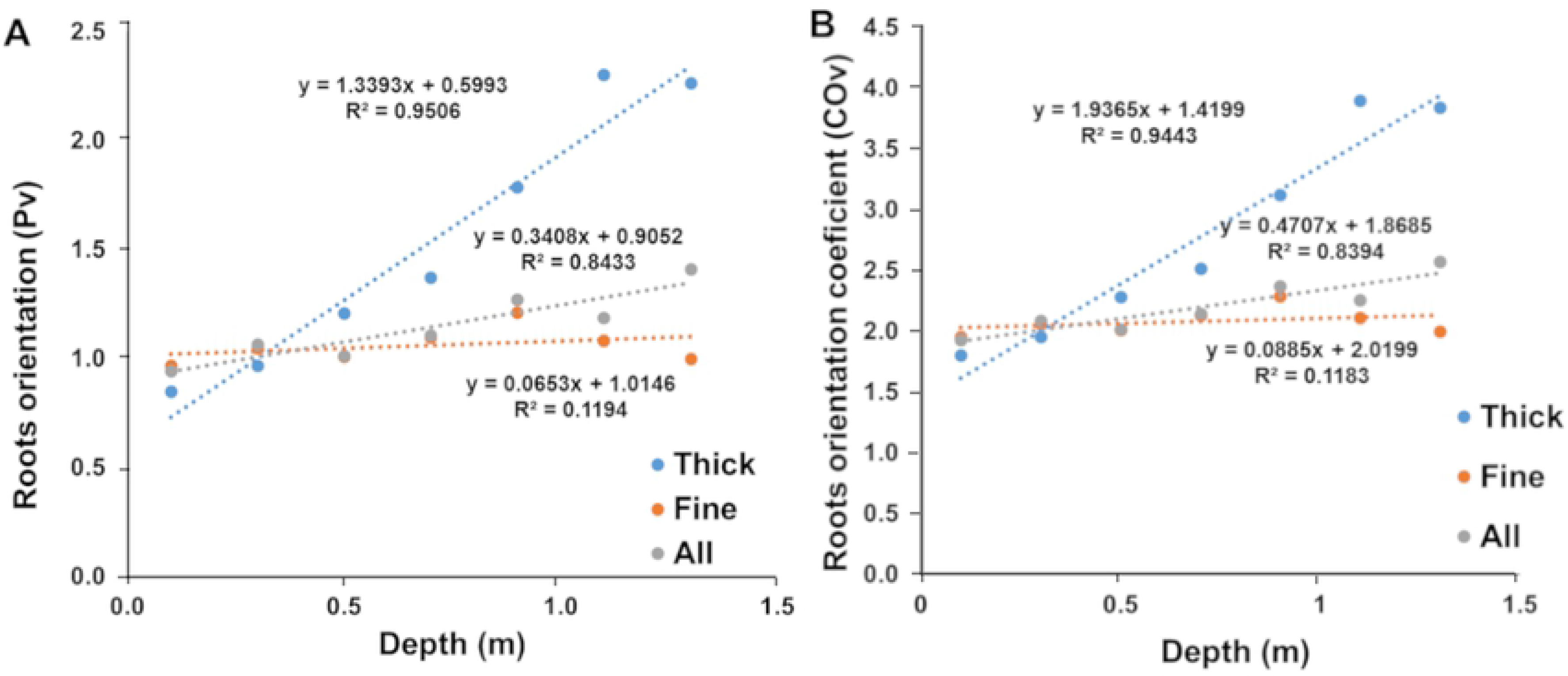
Elaboration of geometric models (all, thick and fine roots). Relationship between soil depth measurements (meters) and the main direction of root growth in relation to a vertical plane (Pv index).

The root direction coefficient of fine roots (COvf) had a low dependence on soil depth (Fig 4A; COvf =0.089 Z+ 2.02; R^2^= 0.118, n= 70). Pf was close to 1 indicating a weak preferential direction of the fine roots. Similarly, COvf had a low dependence on soil depth from 2.02 close to the surface to 2.18 at 1.1 meters depth (Fig 4B). For fine roots, we thus retained a fixed constant value of 2.08 corresponding to the average value of COvf.

For thick roots, root direction (Pvt) and root orientation coefficients (COvt) were strongly dependent on soil depth (Fig 4; COvt= 1.937 Z+1.42; R^2^ = 0.839, n= 70). COvt varied from at 0.10 m depth to almost 4 at 1.3 m depth. This indicates that thick roots tend to grow horizontally close to the surface and that their growth becomes more and more vertical with soil depth.

Altogether, our results indicate that in field conditions pearl millet root orientation depends on soil depth. Thick roots orientation is more sensitive to this than fine roots. We therefore included this information to build 4 models to estimate RLD from RID on the vertical plane taking soil depth (Z: depth in meter) into account:

- an empirical model for all roots: RLDa = 1.83 *RIDa
- a geometrical model for all roots: RLDa = (0.471*Z+1.87)* RIDa
- a geometrical model for fine roots: RLDf= 2.08 * RIDf
- a geometrical model for thick roots: RLDt = (1.937*Z+1.42)* RIDt

### Models validation

Models developed from the data obtained on four varieties during the rainy season 2016 were tested during the dry season 2017 in another field location and with different varieties to maximize the differences between the calibration and validation tests. We used six varieties including five varieties different from those used for model calibration. The quality of the relationships obtained was studied taking into account slope, intercept and regression (R^2^), Nash’s Efficiency Ratio (NE; [22]), root mean square error (RMSE; [23] and mean bias (MB). The results of our statistical tests on the different models are summarized in Table2.

**Table 2.** Models validation analyses.

|  |  |  | Linear regression |  |  |  | Tests |  |  |
| --- | --- | --- | --- | --- | --- | --- | --- | --- | --- |
| Model | Root types | Variety | n | Slope | Intercept | $R^2$ | RMSE (%) | MB (%) | NE |
| Empirical | All | All var | 166 | 1.11 | -1915 | 0.82 | 87 | 24 | 0.58 |
| Geometrical | All | SL423 | 28 | 1.08 | -1259 | 0.83 | 31 | 12 | 0.77 |
|  |  | Souna3 | 28 | 1.06 | -1047 | 0.84 | 26 | 11. | 0.79 |
|  |  | SL28 | 29 | 1.08 | -1253 | 0.87 | 32 | 10 | 0.83 |
|  |  | IBV 8004 | 28 | 1.10 | -1619 | 0.76 | 32 | 14 | 0.70 |
|  |  | GB 8735 | 25 | 1.41 | -3345 | 0.88 | 42 | 11 | 0.78 |
|  |  | ISM 9507 | 28 | 1.07 | -1064 | 0.73 | 36 | 8 | 0.71 |
|  |  | All var | 166 | 1.11 | -1491 | 0.81 | 34 | 11 | 0.77 |
| Geometrical | Fine | SL423 | 28 | 1.12 | -1229 | 0.80 | 35 | 10 | 0.76 |
|  |  | Souna3 | 28 | 1.11 | -1228 | 0.84 | 28 | 11 | 0.80 |
|  |  | SL28 | 29 | 1.14 | -1401 | 0.83 | 37 | 9 | 0.81 |
|  |  | IBV 8004 | 28 | 1.16 | -1754 | 0.78 | 42 | 13 | 0.73 |
|  |  | GB 8735 | 25 | 1.52 | -3464 | 0.89 | 39 | 8 | 0.78 |
|  |  | ISM 9507 | 28 | 1.13 | -1282 | 0.72 | 40 | 7 | 0.70 |
|  |  | All var. | 166 | 1.18 | -1625 | 0.80 | 37 | 10 | 0.76 |
| Geometrical | Thick | SL423 | 23 | 1.38 | -223 | 0.64 | 38 | -0.1 | 0.60 |
|  |  | Souna3 | 26 | 1.59 | -269 | 0.75 | 37 | -13 | 0.61 |
|  |  | SL28 | 26 | 1.21 | -75 | 0.77 | 34 | -9 | 0.73 |
|  |  | IBV 8004 | 21 | 0.77 | 169 | 0.33 | 36 | -7 | 0.28 |
|  |  | GB 8735 | 18 | 0.97 | 60 | 0.32 | 44 | -7 | 0.30 |
|  |  | ISM 9507 | 24 | 0.94 | 165 | 0.51 | 44 | -22 | 0.41 |
|  |  | All var. | 138 | 1.15 | -30 | 0.59 | 39 | -10 | 0.55 |

Characteristics of linear regressions between measured and calculated RLD values. Statistical tests on deviations between measured and calculated RLD values (RMSE: root mean square error), mean bias (MB) and Nash efficiency (NE).

Considering all roots (fine and thick), statistical tests showed that the measured and calculated RLD values were significantly closer with the geometrical model than with the empirical model (Table 2). The MB induced by both models was an underestimation of RLDs for low root intersection densities, generally at depth (Tables 2 & 3, Fig 5A,B).

**Table 3.**
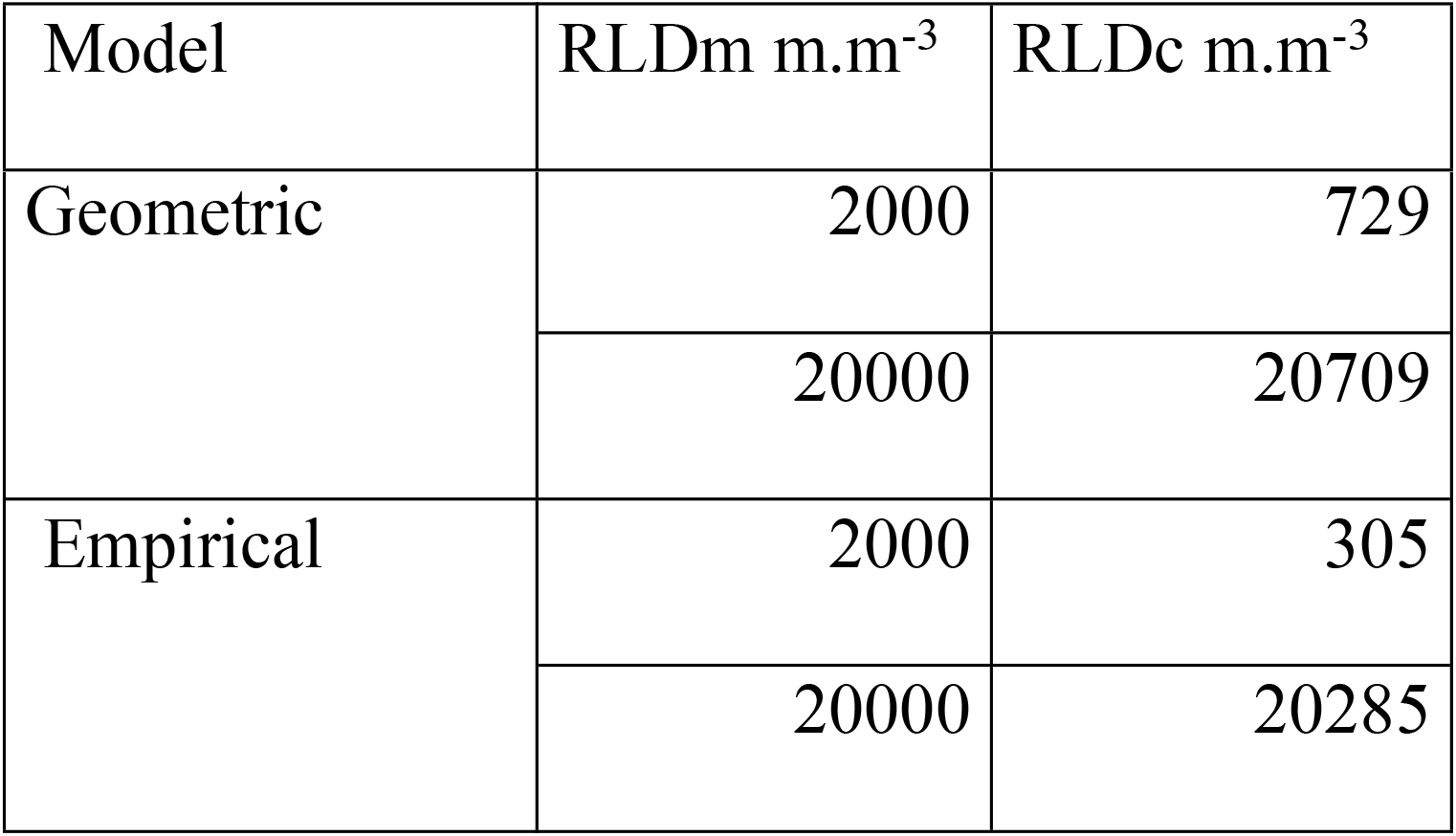
Comparison of the model’s and calculated RLD values of all roots samples using the empirical or geometric models for extreme RLD values.

| Model | RLDm m.m<sup>-3</sup> | RLDc m.m<sup>-3</sup> |
| --- | --- | --- |
| Geometric | 2000 | 729 |
|  | 20000 | 20709 |
| Empirical | 2000 | 305 |
|  | 20000 | 20285 |

**Fig 5.**
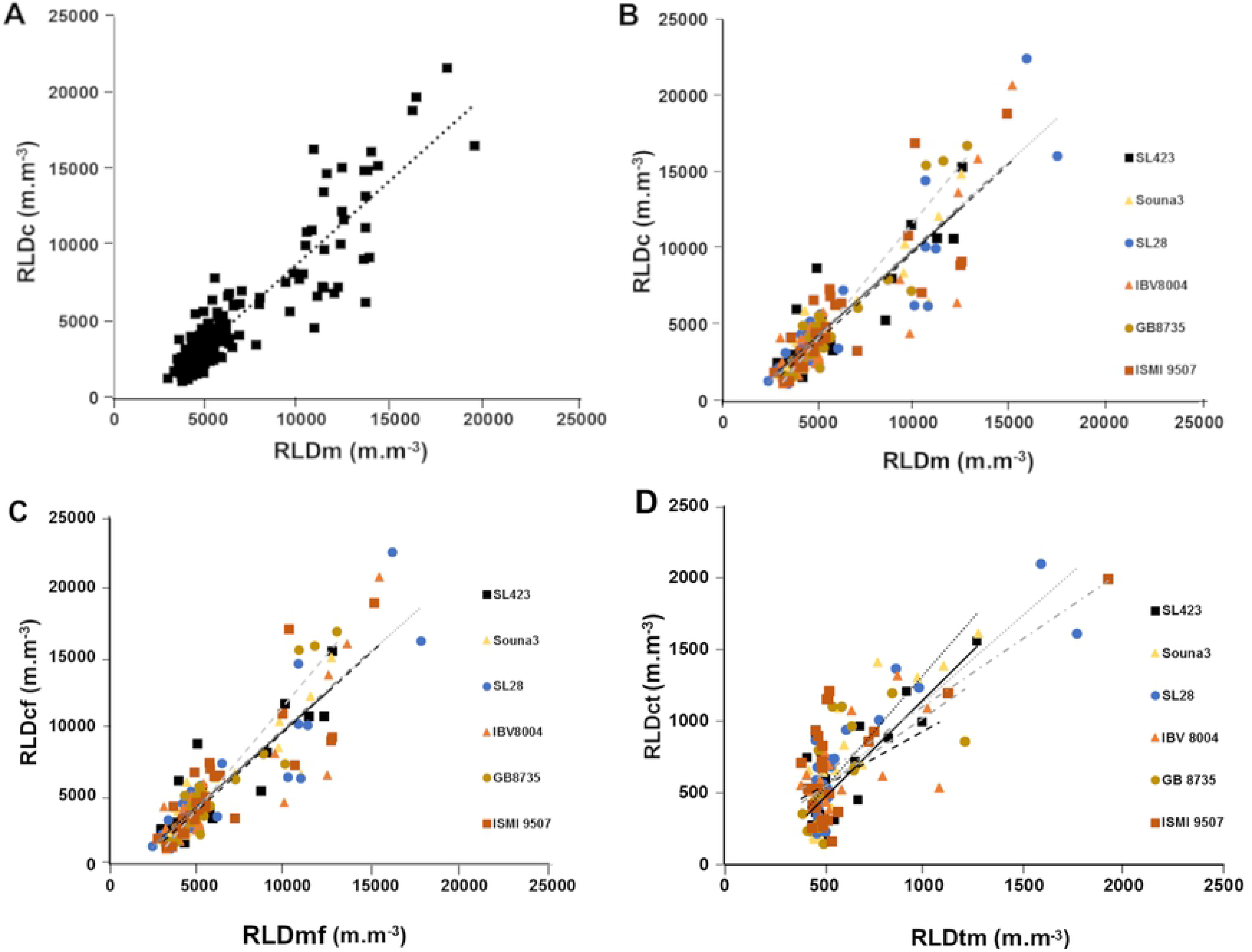
Test of the relationship between measured and calculated RLD for the four proposed models. (A) Empirical model with all varieties bulked, (B) geometric model with all roots, (C) geometric models for fine roots (diameter<1 mm) and (D) thick roots (diameter >1 mm).

The results obtained using models estimating only the fine roots (diameter < 1mm), were close to those obtained using model estimating all roots together. There were good relationships between measured and calculated values of fine roots for each variety except GB 8735 (Fig 5C, Table2).

RLD for thick roots (diameter > 1 mm) ranged from 0 to 2000 m.m^−3^, about ten times lower (Fig 5D) than those for fine roots (Fig 5C). The model construction showed that when the impact density is very low, the relationship between the measured values and those calculated by the thick roots model becomes irregular. It was therefore decided to limit the validation test of the model to a minimum of 70 soil-root intersections per m^2^ (corresponding to at least 3-4 roots on at least one cube face). This value is low, less than one root intersection per dm^2^ (1 per 15 cm square). The validity domain of the “thick root” model will therefore be limited to root intersection counts greater than 70 impacts per m^2^. Despite the removal of the very low RID values, the model does not satisfactorily estimate the RLD when the measured RLD values are low, below 500 m.m^−3^. For these low RLD values, the calculated values were highly fluctuating, but there is no significant bias since the average values per depth of RLD_m_ and RLD_c_ were close (Fig 5D). For higher values of RID and RLD, the model estimated well the RLD averages, although it was still not very accurate when considering sample-by-sample relationships. Among the six tested varieties, two (IBV8004 and GB8735) lead to poorer relationships between the measured RLD values and those estimated by the model (Table 2). These two varieties presented the highest number of samples with RID below 70/m^2^ that were excluded from the analysis and resulted in reduced the number of samples in the dataset. Thus, the percentage of samples eliminated was 25% for IBV8004 and 28% for GB8735 while the percentage of samples eliminated for the other 4 varieties was 12% on average.

Altogether, our experiments validated the following models for RLD estimation from RID:

- for all roots (a): RLDa = (0.471*Z+1.87)* RIDa
- for fine roots (d <1 mm): RLDf= 2.08 * RIDf
- for thick roots (d >1 mm): RLDt = (1.937*Z+1.42)* RIDt

However, the latter was only usable for RLDs > 500 m.m^−3^ and its use is therefore limited.

### Response of pearl millet root system to water stress

We next used our model that takes into account all roots to study the effect of water stress on root architecture in two different pearl millet germplasms, the dual-purpose SL28 variety and the inbred line LCICMB1 (Exp. 3). These two germplasms were grown under irrigated conditions for 40 DAP. Irrigation was then stopped in the drought stress treatment for 31 days while it was maintained in the well-watered treatment. Irrigation was then resumed till the end of the cycle.

Soil water content was followed using Diviner probes and used to compute the fraction of transpirable soil water (FTSW) as previously described [14]. FSTW values below 40% indicate here water-limiting conditions [24]. In the well-watered treatment, FTSW remained above 40% along the soil profile from 30 to 90 DAP (S3 Fig). In the drought stress treatment, water withholding at 31 DAP led to soil drying and induced a reduction in FTSW that fall under 40% between 50 DAP and 70 DAP in the 0-30 cm and 30-60 cm soil layers, respectively (S3 Fig). FTSW was also reduced in the 60-90 cm soil layer reaching 40% at 78 DAP, but remained above 40% below 90 cm throughout the drought stress treatment (S3 Fig). These results are consistent with the ETR / ETM crop ratio values calculated by water balance modeling that estimate ETR / ETM values below 0.3 between 60 and 75 JAP (S2 Fig). Altogether, these results indicate efficient field dry-down and imposition of water limited conditions from topsoil to a depth of around 90 cm in the drought stress treatment.

Agromorphological characteristics were then measured at the end of the cycle (99 DAP). SL28 is a dual-purpose pearl millet variety selected for both fodder and grain production. Accordingly, it shows a very large biomass and grain production compared to the inbred line LCICMB1 in well-watered conditions (Fig 6A,B). Moreover, these two lines showed contrasted responses to drought stress conditions. SL28 showed a very strong and significant reduction in both biomass and grain production in response to water stress while these traits were not significantly affected in LCICMB1 (Fig 6A,B).

**Fig 6.**
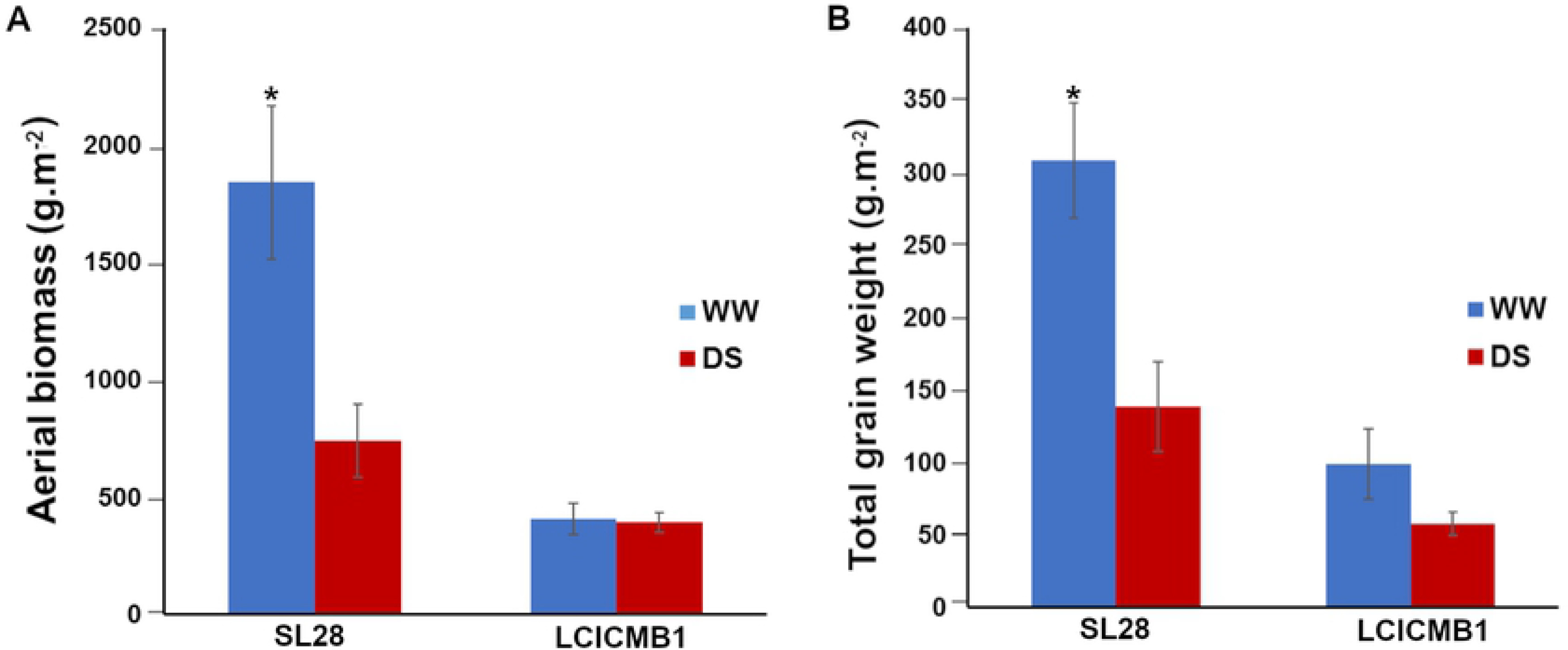
Agromorphological characteristics of SL28 and LCICMB1. (A) aerial biomass (g.m^−2^), and (B) the Total grain weight (g.m^−2^) measured at the end of cycle for WW and DS conditions.

We used the geometrical model for all roots to estimate RLD from RID along soil profiles. Measurements were performed in both well-watered and drought stress conditions for both lines at 43 and 71 DAP, i.e. at the beginning and at the end of the water stress period. Three days after stress application (43 DAP), the RLD profiles were not significantly different for well-watered and drought stress conditions for both lines (Fig 7A,B) indicating that the change in water availability had not significantly impacted root architecture at this stage. However, 31 days after stress application (71 DAP), we observed strong and significant changes in RLD profiles between well-watered and drought stressed plants (Fig 7C,D). For both SL28 and LCICMB1, drought stress led to a significant reduction of RLD in the 0-20 cm soil horizon and to a significant increase in RLD in deep soil layers (> 60 cm; Fig 7C,D). We used the Racine 2.1 application [25] to generate 2D maps of RLD from our data. These maps clearly showed a drastic change in root development occurring both in SL28 and LCICMB1 with a reduction of RLD in topsoil layers and a colonization of deeper soil layer under drought as compared to well-watered conditions (Fig 8A,B). Hence, our data demonstrate that upon drought conditions, both pearl millet lines reduced root growth in the dry topsoil layers and increased their root growth in deeper soil horizons.

**Fig 7.**
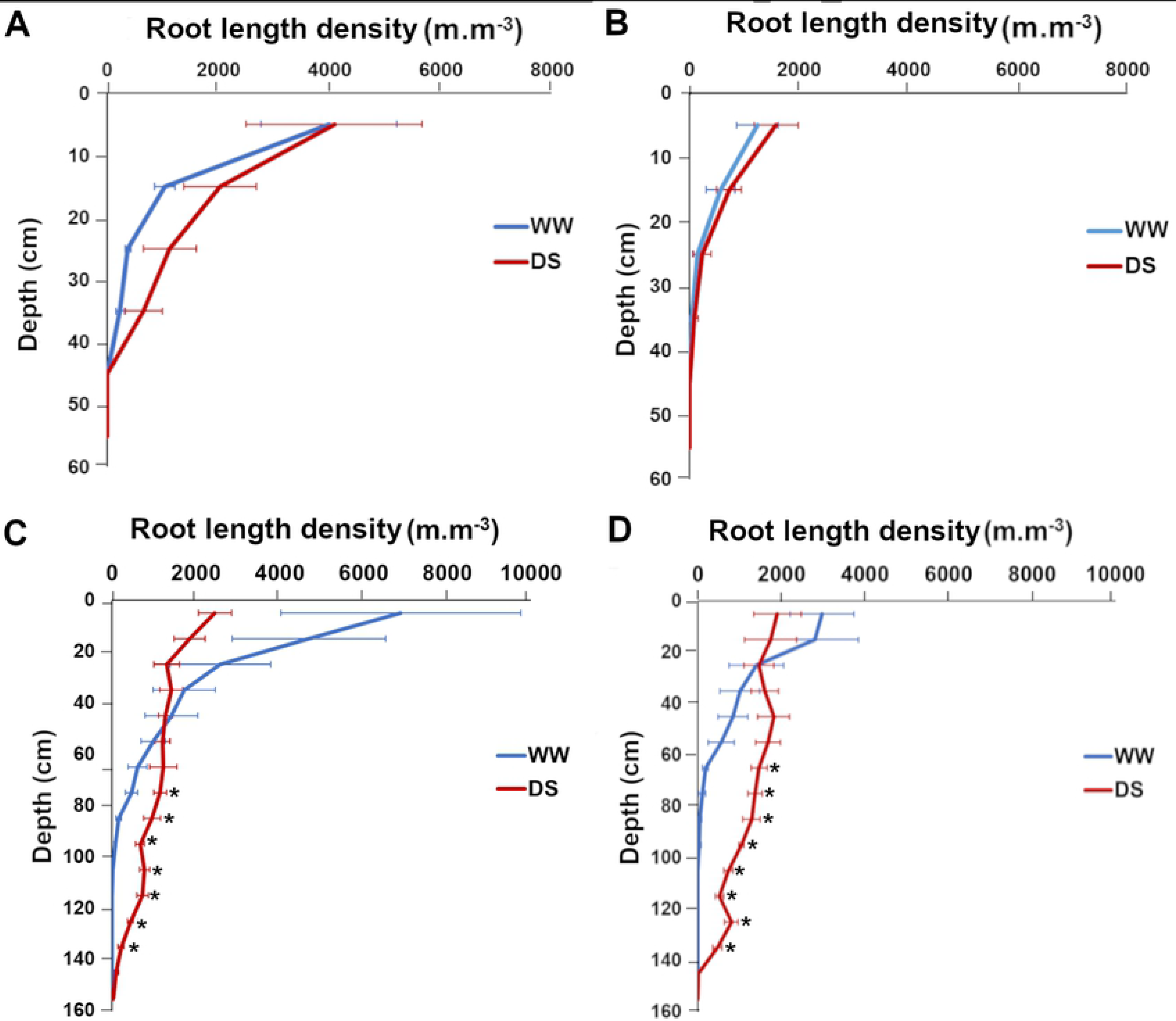
Impact of water deficit on root length density distribution according to depth in SL28 and LCICMB1. SL28 at 43 DAP (A), and 71 DAP (C), LCICMB1 at 44 DAP (B), and 72 DAP (D). Mean of RLD of four plants per variety was considered for each variety.

**Fig 8.**
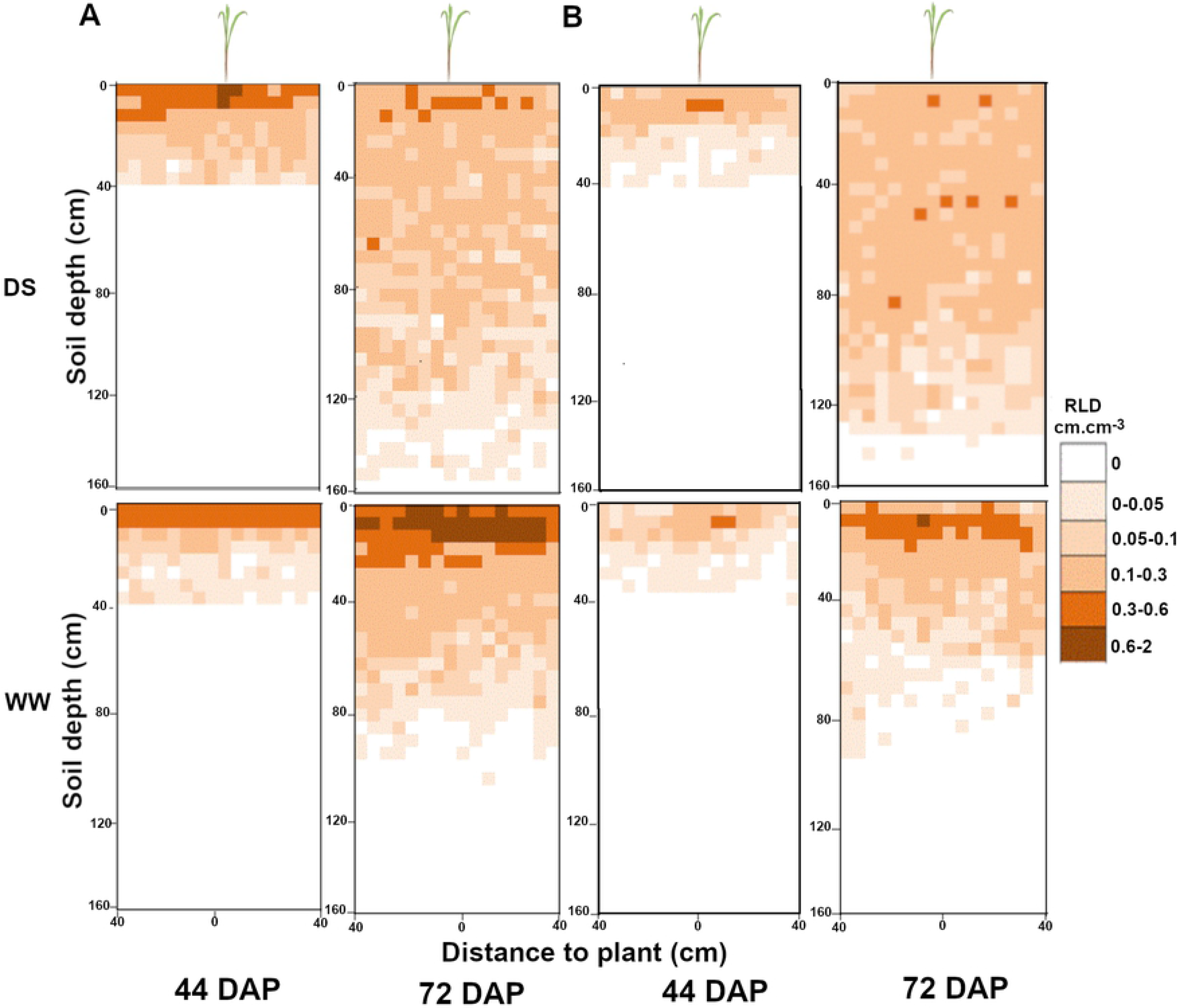
Impact of water deficit on mean root distribution for SL28 and LCICMB1 in the soil profile. Data mapped on a 0.05 × 0.05 m grid like in the field and expressed in root length density (RLD) in WW and DS conditions for SL28 in (A) and the inbred line LCICMB1 in (B).

We used our RLD data to estimate the total length of the root system of SL28 and LCICMB1 per plot surface (m.m^2^) between the soil surface and the root front. Drought stress had contrasted impact on total root length per m^2^ in both lines. We observed a strong and significant increase in total root system length in LCICMB1 and a non-significant reduction in total root length in SL28 (Fig 9A). In the water stress treatment, the ratio between total root length (m.m^−2^) and shoot biomass (g.m^2^) increased in both lines indicating a stronger resource allocation to root growth (Fig 9B). However, this increase was limited and non significant in SL28 while it was large (> 4 times) and significant in LCICMB1 (Fig 9B). Hence, upon drought stress both pearl millet lines seem to reallocate resources to root growth but this reallocation was stronger in LCICMB1.

**Fig 9.**
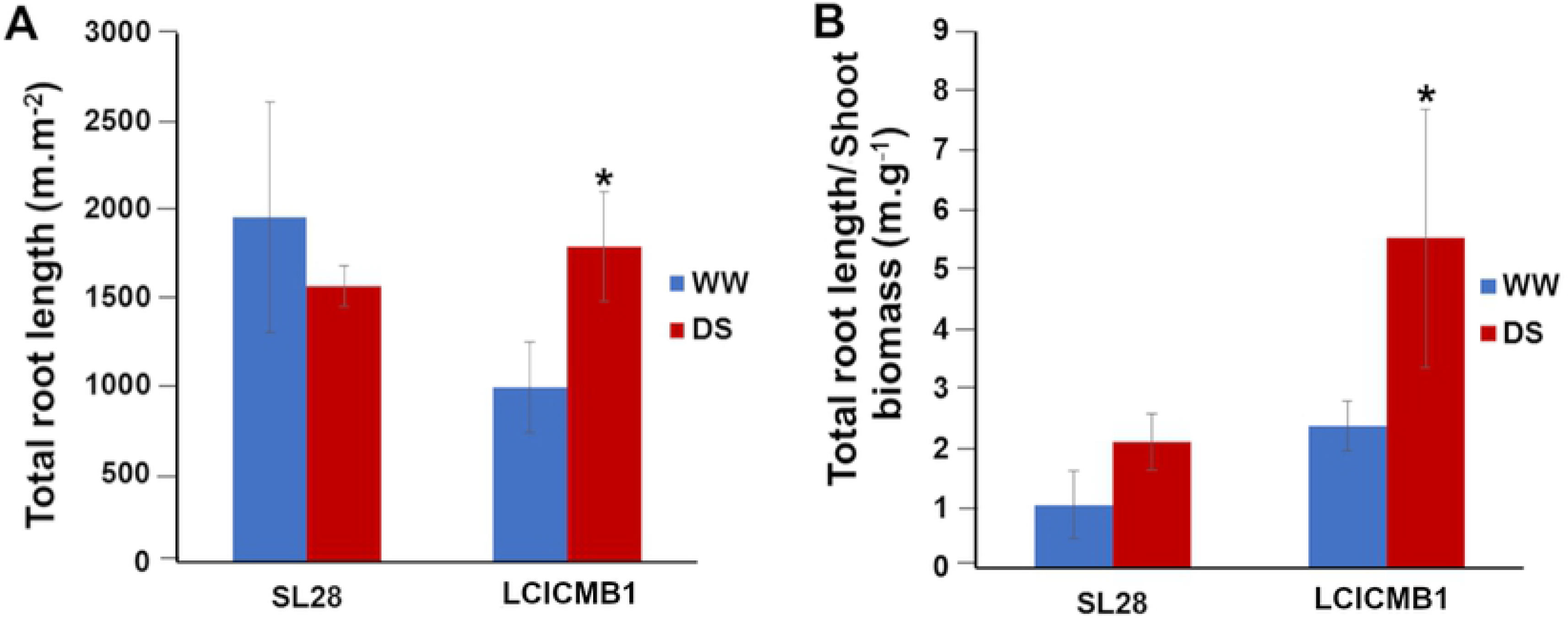
Impact of water deficit on total root length of SL28 and LCICMB1. (A) Total root length (m.m^−3^) measured at 72 DAP at the end of water stress treatment, and (B) ratio between total root length (m.m^−3^) and aerial biomass.

## Discussion

Here, in order to study pearl millet root system in field conditions, we developed a model to evaluate root length density (total length of roots per unit of soil volume; RLD) from root intersection densities (i.e., the number of root impacts on a vertical soil surface; RID). During the development of the model, we observed that pearl millet root growth orientation was dependent on soil depth as already observed for other *Poaceae* species [10, 11]. The dependence was particularly important for thick roots (>1mm diameter) that should correspond either to the seminal or crown roots [13]. The growth of these roots was more or less horizontal in shallow soils and became more and more vertical with increased depth. Conversely, the growth orientation of fine roots, which most likely corresponded to the different types of laterals [13 and 26], was only marginally dependent on soil depth. This led us to develop a model for RLD estimation that considered soil depth as an important variable. This model was validated as the most efficient model to infer RLD from RID. Racine 2.1 [25] was used to manage root data, to calculate RLD and to generate 2D maps of RLD along a soil profile from simple root intersection counts on a vertical plane (trench) thus providing agronomically meaningful information to estimate the efficiency of a root system to acquire water or nutients in different soil horizons. Like most field root phenotyping methods, this method is not high throughput but allows easy and low-cost analysis of root system response to management practices or environmental factors on a reduced sample of accessions. Our results (RLDs and total root length) are consistent with published data obtained in pearl millet using the very labor-intensive but exhaustive monolith method where the root system of a plant is completely dug up by soil layer [27].

As calibrated here, our model will not be suitable for all areas where pearl millet is grown, and in particular to sites with very different soil composition and organization. However, it was developed on a Dior-type of deep sandy soil that is representative of soils where pearl millet is grown in Sahelian West Africa and validated in different fields to ensure it was robust enough. For very different soil types, our model could be simply re-calibrated by measuring the relation between RLD and RID at different soil depth.

Drought is one of the main factors limiting pearl millet yield and drought episodes are predicted to increase in number and length in the future in West Africa [6, 28]. Previous studies suggested that pearl millet tolerance to dry environments could be due to mechanisms regulating water use efficiency and limiting water loss rather than to improved water acquisition [29]. Interestingly, an expansion of gene families involved in cutin, suberin and wax biosynthesis was observed in pearl millet compared to other cereals and a potential QTL for biomass production under drought was found to co-locate with a gene encoding 3-ketoacyl-CoA synthase that catalyzes the elongation of C24 fatty acids during both wax and suberin biosynthesis [14] thus supporting the link between transpiration barriers and drought resistance in pearl millet. Experiments using lysimeters indicated that temporal patterns of water use, rather than total water uptake, were essential for explaining the terminal drought tolerance of pearl millet genotypes containing a terminal drought tolerance QTL [30]. Therefore, this terminal drought QTL did not affect the water extraction capacity of the root system. Moreover, it was reported that water stress did not lead to increased water uptake from deep soil suggesting that drought did not lead to a deeper root system [29]. However, the corresponding experiments were performed in pots or lysimeters that limit the full expression of root architecture component compared to field conditions.

We therefore used our phenotyping method to analyze the response of pearl millet root system to water stress during the vegetative phase in field conditions. Our experiments were performed during the dry season on two germplasms with contrasted characteristics: a dual-purpose variety that develops a large aerial biomass and is sensitive to drought and an inbred line with a more limited biomass and that is less sensitive to drought. Our results clearly show that water stress leads to a reallocation of carbon for root growth combined to a reduction of RLD in topsoil layers and to an increase in root system depth. It demonstrates that upon drought stress, pearl millet increases its root growth in deeper soil layer that retain some water. While we cannot conclude from such a small sample, we can hypothesize that this response is adaptive, i.e., that it contributes, with other strategies such as reduction in water loss and temporal regulation of water uptake, to pearl millet tolerance to drought stress. Further work will be needed to test this hypothesis.

In conclusion, we developed a simple way to evaluate and map pearl millet RLD distribution in field conditions. This opens the perspective to characterize the impact of a number of environmental factors and management practices on field-grown pearl millet root system development.

## Acknowledgements

This research was supported by the Sustainable Intensification Innovation Lab project (SIIL / Feed the Future) through the United States Agency for International Development (USAID, grant 929773554). A. Faye was supported by a PhD grant from the SIIL. We thank V. Vadez (IRD, France) and M.J. Bennett (University of Nottingham, UK) for critical reading of our manuscript.

## Supporting information

**S1 Fig. Climatic data for Exp. 1 & 2**

**S2 Fig. Climatic data for Exp. 3**

**S3 Fig. Soil water content during Exp. 3**

**S1 Table. Student t-test on the effect of different factors on the preferential orientation indices (P) of fine (P_f_), thick (P_t_) and all roots (P_a_)**

**Supporting Data. Data from all the experiments.**

